# DNA extrusion size determines pathway choice during CAG repeat expansion

**DOI:** 10.1101/2025.05.12.653571

**Authors:** Mayuri Bhatia, Ashutosh S. Phadte, Anna Lakhina, Anthony R. Monte Carlo, Sarah Barndt, Anna Pluciennik

## Abstract

DNA triplet repeat expansion is the mutational cause of neurodegenerative disorders such as Huntington’s disease, myotonic dystrophy type 1, and fragile-X related disorders. There is a general consensus that recognition of extrahelical extrusions or hairpin-loop structures (formed by strand slippage) by the DNA mismatch repair protein MutSβ leads to repeat expansion by a mutagenic repair process. By contrast, the FAN1 nuclease prevents triplet repeat expansion, the molecular basis of which was explained by our recent finding that FAN1 nuclease cleaves and initiates removal of extrahelical extrusions. We have proposed that competition for extrusion binding between FAN1 and MutSβ governs the outcome of the opposing effects of these two pathways. Here we show that extrusions containing 2-3 triplet repeats are recognized and processed by both FAN1 and MutSβ pathways. However, a single triplet extrusion escapes FAN1 cleavage and is exclusively processed by MutSβ-dependent MMR, leading to repeat expansion. Thus, the size of the extrahelical extrusions formed by strand slippage events affects the ultimate fate of the repeat elements, and controls the bias between repeat expansion or stability. These findings provide new insights into the role of DNA structural dynamics in establishing pathway choice in DNA repair.

## INTRODUCTION

DNA repeats are found throughout the genome, and are polymorphic in length due to their genetic instability (1). These repeat length changes are triggered by the formation of transient unusual DNA structures (extrahelical extrusions) during DNA metabolic processes (2–4). The detrimental consequences of repeat instability are exemplified by triplet repeat expansions that cause a number of neurodegenerative diseases such as Huntington’s disease (HD), myotonic dystrophy type1 (DM1), and fragile-X related disorders (FXDs) (5,6). Genome-wide association studies (GWAS) in HD patients have recently identified several genetic modifiers of disease onset age (7–15) that map to DNA repair pathways. These include *FAN1* (16) and the DNA mismatch repair (MMR) genes *MSH3*, *PMS1*, *PMS2* and *MLH1* (reviewed in (17–19)). Of these, MSH3, MLH1, MLH3, and PMS1 have been implicated as drivers of somatic triplet repeat expansion (11,20–32), a counterintuitive finding in light of the critical function of the MMR pathway in mutation avoidance. By contrast, the FAN1 nuclease attenuates repeat expansion (12,32–34). Thus, the FAN1 and MMR pathways exert opposing effects on repeat expansion. However, the molecular mechanisms that govern which pathway dominates are not well understood.

There is a broad consensus that the mutagenic role of MMR is triggered by recognition of triplet repeat extrahelical extrusions by the MutSβ heterodimer (MSH2/MSH3) (19,32,35), a protein known to be involved in rectification of insertion/deletion loops of 2-10 nucleotides (36–38). Repeat expansion also requires the MutL homologs MLH1, MLH3, and PMS1, suggesting a role for the MutLγ (MLH1/MLH3) and MutLβ (MLH1/PMS1) heterodimers in this process. Recently, we and others have shown that extrahelical extrusions can also be recognized by the FAN1 nuclease (39,40), an enzyme previously known primarily for its role in repair of DNA interstrand crosslinks (41–45). FAN1 nucleolytic cleavage of CAG and CTG extrusions requires DNA-loaded PCNA, resulting in the removal of such structures by a short patch excision/repair process (39). FAN1 action in this manner is highly strand-specific, with activity restricted by the orientation of DNA-loaded PCNA to the strand that contains an extrusion. This reaction requires a physical interaction between FAN1 and PCNA, the nature of which has been revealed by cryo-EM structures of the FAN1-PCNA-DNA ternary complex (46,47). Disruption of the FAN1-PCNA interaction by peptide inhibitors or by mutation of the PCNA-binding interface of FAN1 results in abrogation of PCNA-activated FAN1 cleavage of extrahelical extrusions.

As noted above, triplet repeat expansion is a net outcome of opposing interplay between the FAN1 and MMR pathways. It has been suggested that a physical interaction between FAN1 and MLH1 may inhibit CAG repeat expansion by sequestering MLH1 heterodimers, thereby depriving the MutSβ-dependent MMR machinery of a key catalytic component (48,49). Based on biochemical findings, an alternative (and mutually non-exclusive) model posits that competition between FAN1 and MutSβ for CAG extrahelical extrusions might determine this pathway choice (39,50). It is therefore plausible that relative protein levels of FAN1 and MutSβ at the relevant DNA loci may dictate which pathway dominates.

Because DNA strand-slippage and misalignment within triplet repeat tracts can result in extrahelical extrusions of varying sizes, we evaluated the effects of extrusion size on FAN1 nuclease activity. We show that FAN1 nuclease displays a strong preference for cleaving extrusions of 2-3 repeats over a single triplet extrusion. This effect correlates with the propensity of the extrusion to induce a kink in the DNA, with a bend angle of >90° being conducive for FAN1 cleavage. By contrast, MutSβ shows no preference for (CAG)_2, 3_ over (CAG)_1_, and supports MutLα incision on these substrates with comparable facility. Our observations imply that (CAG)_1_ extrusions within triplet repeats exclusively provoke MutSβ-mediated MMR, and escape removal by FAN1, thereby leading to expansions.

## MATERIALS AND METHODS

### Proteins and DNA substrates

Human recombinant FAN1 or its catalytic mutant (D960A) were purified from *E. coli* cells as described previously (39). Recombinant human proliferating cell nuclear antigen (PCNA) was purified from *E. coli* harboring plasmid pET11a-PCNA as described (46). Human replication factor C (RFC), MutLα, and MutSβ were purified from baculovirus infected Sf9 cells according to published protocols (37,51,52).

The DNA substrates harboring (CAG)_1, 2, or 3_ extrahelical extrusions were prepared as described (35), using phagemid constructs that harbor sequences bounding the CTG repeat within the 3′-untranslated region of the human DMPK gene (19 nt 5′ and 40 nt 3′). Oligonucleotide-based DNA substrates were prepared by annealing of HPLC-purified complementary oligonucleotides (Integrated DNA Technologies) (Supplementary Table 1). The annealed substrates were run on 10% polyacrylamide gels in 1X TBE buffer to confirm the annealing efficiency. The double-stranded substrates were then radiolabeled using [α-^32^P] d(GTP) and exo^-^ Klenow at the 3′ end on the extrusion harboring strand (see Fig. 1). For the 5′ radiolabeled DNA substrates, DNA oligonucleotides were labeled with [γ-P^32^] (ATP) and T4 polynucleotide kinase, followed by annealing with appropriate complementary strands to form double stranded DNAs harboring an extrusion or homoduplex controls. All oligonucleotide sequences used in this study are detailed in Supplementary Tables 1-3.

**Figure 1.**
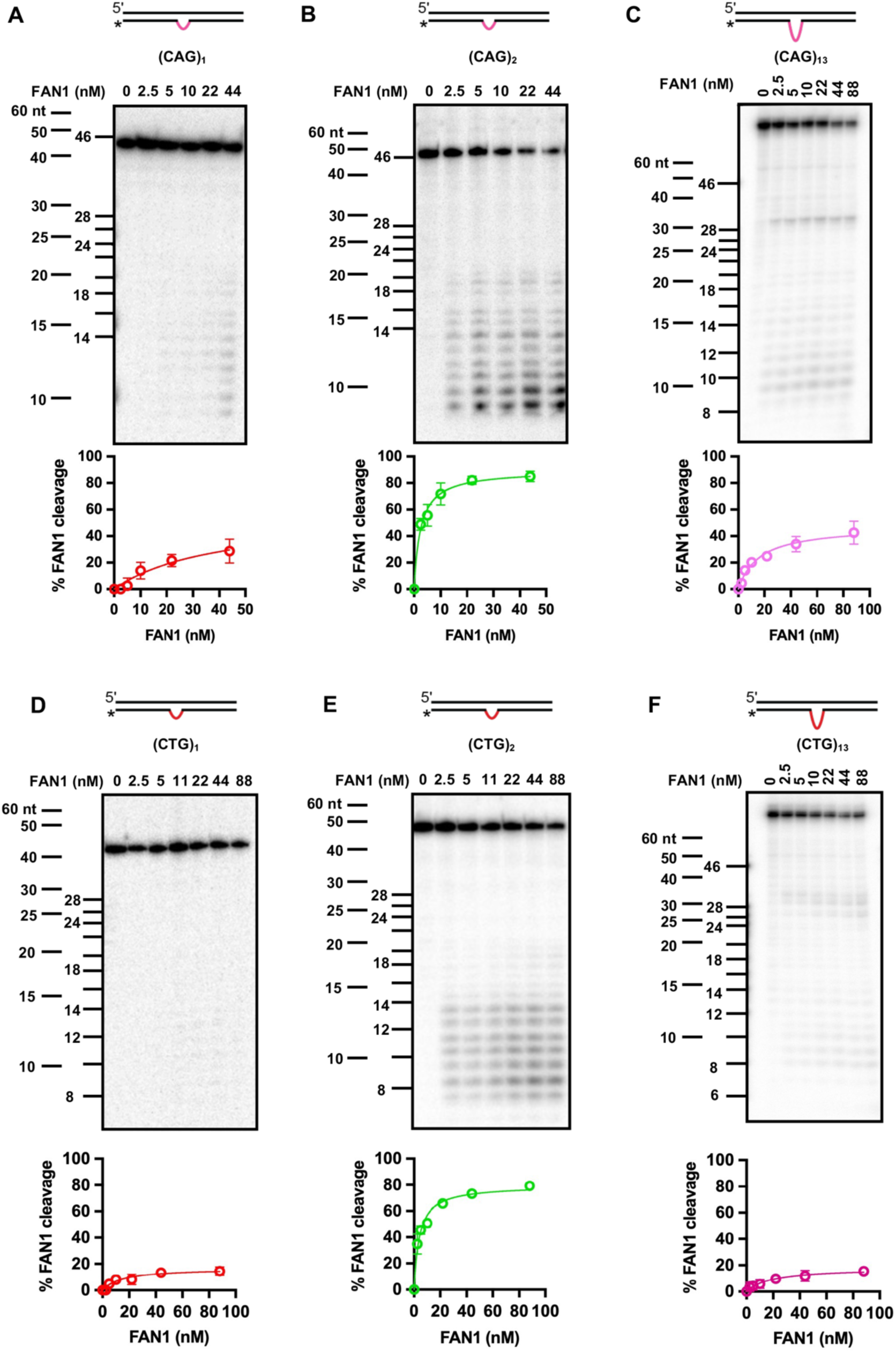
FAN1 preferentially cleaves extrahelical extrusions containing 6-9 nucleotides. Five nanomolar 3’-end radiolabeled DNA substrates harboring extrusions of (**A**) (CAG)_1_, (**B**) (CAG)_2_, (**C**) (CAG)_13_, (**D**) (CTG)_1_, (**E**) (CTG)_2_, (**F**) (CTG)_13_ were incubated with increasing concentrations of FAN1 for 10 min at 37 °C in the buffer containing 5 mM MgCl_2_ and 70 mM KCl (see Materials and Methods). The samples were collected at different time points and resolved on 20% denaturing PAGE. Representative images are shown. Two DNA markers were used (i) marker of the same sequence as the DNA substrate (see Supplementary Table 2), and (ii) commercial 60-10 nucleotide (nt) marker. Please note that for (CAG)_1_ and (CAG)_2,_ the highest concentration of FAN1 is 44 nM and for the remaining substrates FAN1 concentration is 88 nM. Quantification of percent of FAN1 cleavage is shown below each respective image. Graphs represent mean values ± SD of at least 3 independent experiments. Also see Supplementary Figure 1.

### FAN1 nuclease assays on end-labeled linear DNA substrates

Reactions (20 μL) contained 5 nM of 3’-radiolabeled DNA substrate harboring (CAG)_1, 2, 3, 4, 5, or 13_ or (CTG)_1, 2, 3, 4, 5, or 13_ extrahelical extrusion (or a homoduplex control) and FAN1 (0 – 88 nM, or as indicated) in 25 mM HEPES-KOH pH 7.5, 5 mM MgCl_2_, 0.05 mg/mL BSA (Sigma Aldrich, cat # 10711454001) and 70 mM KCl. The reaction mixes were incubated at 37 ⁰C for 10 minutes for all titration assays. Time-dependent assays were done in the same manner by incubating 5 nM DNA substrate with 11 nM FAN1 and collecting samples at indicated time points. The reactions were terminated by addition of formamide to a final concentration of 70%. The samples were incubated at 95 °C for 5 minutes and resolved on 20% PAGE containing 8 M Urea, followed by phosphorimaging. The bands were quantified using ImageJ software. Percent FAN1 nuclease activity was calculated as a ratio of cleavage products to the total intensity in the lane. In order to evaluate the FAN1 cleavage products, sequence-specific markers were used for CAG/CTG substrates (sequence identical to the strand harboring extrusion (see Supplementary Table 2)). Commercial DNA marker (10-60 nt and 20-100 nt) was also used (cat # 51-05-15-01 and 51-05-15-02).

To evaluate the FAN1 processivity, 10 nM of 3’-radiolabeled (CAG)_2_ DNA substrate was incubated at 37 ⁰C with 2.5 nM FAN1 nuclease in buffer composed of 25 mM HEPES-KOH pH 7.5, 5 mM MgCl_2_, 0.05 mg/mL BSA, and 70 mM KCl. After 10 seconds of incubation, 250 nM of 3’-Cy3-labelled (CAG)_2_ DNA competitor or buffer were added and incubated at 37 ⁰C for the indicated time. Samples were collected and analyzed as above.

### FAN1 and MutLα endonuclease assays on circular DNA substrates

In order to assess how extrusion size influences FAN1 and MutLα nuclease activity, reactions containing components conducive for FAN1 and MutLα nuclease activity were carried out in 20 mM Tris pH 7.6, 125 mM KCl, 1.5 mM ATP, 1 mM glutathione, 5 mM MgCl_2_ and 2.5% glycerol. To assess for MutLα endonuclease activity, 2.5 nM of the double-stranded circular DNA substrate harboring either (CAG)_1_, (CAG)_2_ or (CAG)_3_ extrahelical extrusion was incubated in the presence of either 10.5 nM RFC, 36.5 nM PCNA, 43 nM MutSβ (MSH2-MSH3 heterodimer) and 7 nM MutLα (MLH1-PMS2 heterodimer) as indicated. Reactions were carried out at 37 °C for 15 minutes, after which they were terminated by the addition of 6X alkaline gel loading dye – 300 mM NaOH, 6 mM EDTA, 18% Ficoll 400, Bromocresol green, to a final concentration of 1X. Reaction mixes were vortexed and allowed to incubate for 10 minutes at room temperature.

To assess FAN1 nuclease activity, 2.5 nM of the substrate containing either (CAG)_1_, (CAG)_2_ or (CAG)_3_ extrusion was incubated in the presence of 10.5 nM RFC, 36.5 nM PCNA and 2.5 nM FAN1 as indicated. Reaction mixes were incubated at 37 °C for 10 minutes, after which they were terminated by the addition of 6X alkaline loading dye to a final concentration of 1X as above. Samples were run on 1% alkaline agarose gels in 50 mM NaOH, 1 mM EDTA at 28 V for 450 Vh. The gels were dried, rehydrated and subjected to Southern blotting using ^32^P-labeled oligonucleotide probes targeting either the extrusion containing strand (Fwd 2943), or the complementary strand (Rev 2970). The gels were exposed to phosphorimager screens, followed by quantitation using a Molecular Dynamics Phosphorimager.

### Extrusion-dependent electrophoretic mobility

DNA substrates harboring (C), (CA), (CAG)_1_, (CAG)_2_, (CAG)_3_ extrusions at different positions of the DNA duplex (Supplementary Table 3) were mixed with Ficoll (to 14% final concentration) and resolved through 15% polyacrylamide gel for 10 hours. The gel was then stained with 0.01 mg/ml ethidium bromide for 20 minutes followed by washing with deionized water for 40 minutes and imaged using Bio-Rad chemidoc imaging system. The migration distance was calculated by measuring the distance from the respective wells for each substrate in ImageJ software. The distance was plotted against each substrate to evaluate the variability in substrate migration due to positioning and size of extrusion present on the heteroduplex.

### Surface plasmon resonance analysis

FAN1-DNA interaction was determined by surface plasmon resonance spectroscopy (SPRS) using BIAcore 3000. The DNA substrates were prepared by annealing of oligonucleotides (Supplementary Table 1) to yield DNA substrates harboring (CAG)_1, 2, 3, 4, 5, or 13_ or (CTG) _1, 2, 3, 4, 5, or 13_ extrahelical extrusions or homoduplex control. These DNAs contained a 5’-terminal biotin on the DNA strand harboring the extrusion and a 5’-terminal digoxigenin on the opposite strand. The streptavidin chip (Xantec, cat # SAHC200M) was prepared as per manufacturer instructions and derivatized with approximately 150 – 200 response units (RUs) of biotinylated heteroduplex or homoduplex DNA substrates. The flow cell 1 was considered as a reference, the control homoduplex was immobilized on flow cell 2, and the appropriate heteroduplex on flow cell 3. The free end of the DNA substrates were blocked by flowing 50 nM anti-digoxigenin antibody (Roche, cat #11333089001) over flow cells 2 and 3. FAN1 was injected at a flow rate of 20 μL/min in a buffer containing 20 mM Tris-HCl (pH 7.6), 70 mM NaCl, 10 mM CaCl_2_, 1 mM EDTA, 0.01% P-20, 1 mM DTT, and 0.05 mg/ mL of BSA. Experiments were performed at 5 °C. Dissociation constants for FAN1-DNA interactions were estimated using SPRS by titrating FAN1 (5, 10, 25, 50, 100, 200 and 400 nM) over the chip-bound DNA. The chip was regenerated by flowing 1 – 3 M NaCl after each kinetic cycle. The data is reported in form of time-dependent response sensograms. The steady state affinity curve was plotted from the maximum response attained 5 seconds before the end of injection for each FAN1 concentration flown over the bound DNA on the chip to study time-independent stability of the complex. One-site binding kinetic model, Hyperbola, was used in Graphpad Prism 9.0 software for affinity plot to calculate the dissociation constant (K_d_), representing the concentration of the analyte (protein) at which half of the binding sites available on the immobilized ligand (DNA) are occupied for all CAG substrates and respective homoduplexes.

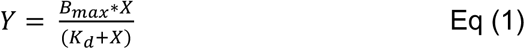

Where Y is the maximum response achieved for each flowing concentration of protein, X is protein concentration and B_max_ is the maximum binding capacity.

MutSβ-DNA binding studies were performed using Biacore 1K. DNA substrates with (CAG)_1_, (CAG)_2_, (CAG)_3_ or (CAG)_13_ extrusion or a homoduplex control were immobilized on a 6-flow cell chip, where, flow cells 1, 3 and 5 were used as a reference against the flow cells 2, 4 and 6 as active surfaces and the biotinylated heteroduplex and homoduplex controls were immobilized. The chip surface was activated by 1M NaCl / 50 mM NaOH conditioning followed by DNA substrate immobilization on flow cells 2, 4, and 6 at 10 μL/min with a target level of 15-25 response units. Immobilization and the binding assays were done in buffer containing, 10 mM Tris-HCl pH 7.6, 150 mM NaCl, 10 mM MgCl_2_, 1 mM EDTA, 1mM dithiothreitol (DTT), 0.005% P-20. Increasing concentrations of MutSβ in the running buffer (0.1 to 100 nM) were stored in the sample chamber at 10 °C until injection. The multicycle kinetic method was used with 210 sec injection of each concentration of MutSβ at 20 μL/min, followed by a 120 sec dissociation time. The chip was regenerated by washing with 1 mM ATP in the running buffer after each injection.

A 1:1 binding model was used for kinetic fitting to the sensograms for each MutSβ concentration using Biacore 1K evaluation software to study the time-dependent interaction between protein and DNA. The rate of MutSβ-DNA interaction was evaluated in the presence and absence of ATP. The binding rate constants for association and dissociation were evaluated using the following equations:

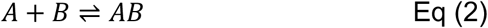

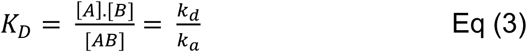

Where K_D_ is the equilibrium dissociation constant, A is the analyte (protein), B is the ligand (immobilized biotinylated DNA), and k_a_ and k_d_, representing association and dissociation, respectively. The association vs. dissociation rates for (CAG)_1,2,3,13_ and homoduplex substrates were tested for statistical significance using one-way ANOVA. Further, to study time-independent interaction of MutSβ and DNA, steady-state affinity model was fitted using the Biacore 1K evaluation software for the maximum response attained, 5 seconds before the injection ends, for each MutSβ concentration. The time-independent response vs MutSβ concentration plot was used to determine the K_D_ value for each DNA substrate:

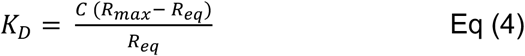

Where, C is the concentration, R_max_ is the maximum feasible response achieved by interaction between A and B, R_eq_ is the response attained from interaction between DNA and each concentration of protein at equilibrium.

## RESULTS

### FAN1 preferentially cleaves extrahelical extrusions containing 6-9 nucleotides

DNA extrahelical extrusions are formed within repetitive sequences via slippage of the two complementary strands during DNA metabolic processes (53–55). Our recent studies have uncovered a novel role for FAN1 in limiting triplet repeat expansion by removal of (CAG)_2_ or (CTG)_2_ extrahelical extrusions that form within CAG/CTG repeat tracts associated with several neurodegenerative diseases (39). The nuclease activity of FAN1 is provoked by such extrahelical extrusions, as judged by the inertness of the enzyme on homoduplex DNAs (39). However, structural features of the extrusions (such as size or sequence composition) that govern FAN1 nuclease function on such DNAs are not well understood. Therefore, in order to evaluate the effect of the extrusion size within the DNA substrate on FAN1 nuclease activity, we constructed DNA substrates with varying numbers of CAG or CTG repeats forming the extrusion (1, 2, 3, 4, 5, and 13 CAG-repeats within the extrusion as shown in Supplementary Table 1). The DNA substrates were radiolabeled at the 3’ end and evaluated in FAN1 nuclease assays as described (39). We observed that DNA substrates harboring (CAG)_2_ or (CAG)_3_ extrusions are cleaved efficiently even at low nanomolar concentrations of FAN1 (Figure 1B, and Supplementary Figure 1A), whereas a catalytic mutant of FAN1 (D960A) (43,45) purified using the same method does not display detectable activity (Supplementary Figure 2A, B). In comparison, (CAG)_1_ extrusions are poor substrates for FAN1, with 5 to 10-fold higher amounts of the enzyme required to achieve similar levels of cleavage of (CAG)_1_ relative to (CAG)_2_ or (CAG)_3_ (Figure 1, and Supplementary Figure 1). This effect is even more striking on DNAs harboring (CTG) extrusions, which require > 35-fold higher amount of FAN1 to achieve cleavage of (CTG)_1_ comparable in extent to (CTG)_2_ or (CTG)_3_ (Figure 1D, E, and Supplementary Figure 1D). Interestingly, FAN1 displays substantially lower levels of activity on (CAG)_13_ or (CTG)_13_ relative to extrusions containing two repeats (Figure 1C, F). These observations indicate that the FAN1 nuclease activity is exquisitely sensitive to the size of the extrusion.

### FAN1 cleavage rates depend on the size of the extrusion formed by triplet repeats

In order to further evaluate the kinetics of FAN1 nuclease on these DNA substrates, we performed time course experiments (Figure 2A-C and Supplementary Figures 3, 4). The rates of initiation of FAN1 cleavage under these experimental conditions displayed a ∼2.5-fold preference for (CAG)_2_ (0.005 s^-1^) in comparison to (CAG)_3_,_5_ (∼0.002 s^-1^); and a ∼5-fold preference in comparison to (CAG)_4_,_13_ (0.001 s^-1^). DNAs harboring a (CAG)_1_ extrusion are poor substrates for FAN1 nuclease: cleavage is below detection limits within the first minute of the reaction.

**Figure 2.**
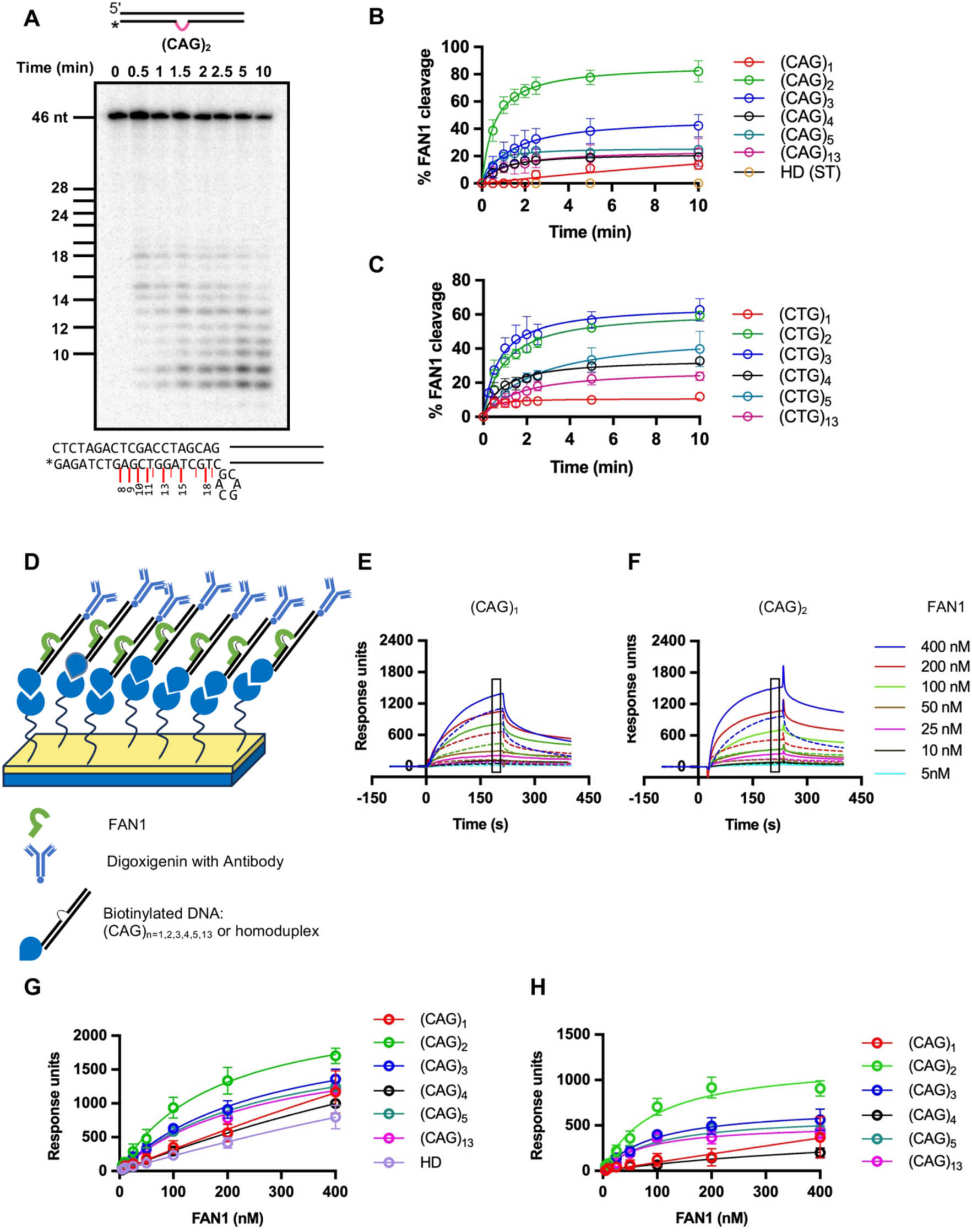
FAN1 cleavage rates depend on the size of the extrusion formed by triplet repeats. (**A**) Five nanomolar 3’-end radiolabeled (CAG)_2_ extrusion harboring DNA substrate was incubated with 11 nM FAN1 at 37 ⁰C in the buffer containing 5 mM MgCl_2_ and 70 mM KCl. The samples were collected at different time points and resolved on 20% denaturing PAGE. The DNA marker is of the same sequence as the DNA substrate used in the experiment. Major FAN1 cleavage sites are depicted by red lines on the DNA substrate sequence. (**B**) Quantification of FAN1 cleavage on DNA substrates harboring (CAG)_1, 2, 3, 4, 5, or 13_ extrahelical extrusions based on experiments as shown in (A) and Supplementary Figure 3. Graph represents mean values ± SD of n=3 independent experiments (except for (CAG)_4_ where data is based on 2 independent experiments). (**C**) Quantification of FAN1 cleavage on DNA substrates harboring (CTG) _1, 2, 3, 4, 5, or 13_ extrusions based on experiments as shown in Supplementary Figure 4. Graphs are presented as mean values ± SD of n=3 independent experiments (except for (CTG)_3_ data is based on n=2 independent experiments). FAN1 binding is driven by the extrusion size. (**D**) Schematics of the experimental design for the surface plasmon resonance experiments. (**E**) The assembly of FAN1-DNA binary complex was scored by SPRS using a sensor chip derivatized with 40-bp DNA substrate harboring (CAG)_1_ or (CAG)_2_ extrusion (solid lines) or (**F**) 46-bp homoduplex control (dashed lines). Sensograms show mass response units upon flow of solutions containing increasing concentrations of FAN1 (as indicated). (**G**) Apparent affinity of FAN1 for different DNA substrates was determined from SPRS experiments like those described in (**E, F**). The data were fit to a hyperbola using Eq (1) (Experimental Procedures). We observed a substantial FAN1 binding to homoduplex control, although the binding curves do not saturate. Data are based on n=3 for (CAG)_2_, n=2 for (CAG)_1,3,13_ independent experiments with error bars representing SD. Data for (CAG)_4,5_ are based on a single enzyme titration experiment. (**H**) Data from (**G**) were also plotted after subtraction of homoduplex values from those obtained with (CAG) extrusion DNA substrates in order to correct heteroduplex binding for nonspecific effects.

The differences between the rates of enzymatic cleavage may be attributed at least in part to the binding preference of FAN1 for these substrates. Surface plasmon resonance experiments (SPRS) clearly established that FAN1 preferentially binds (CAG)_2_ over all other substrates, with limited binding observed for (CAG)_1_ (Figure 2D-H, and Supplementary Table 4). It should be noted that substantial levels of binding of FAN1 to homoduplex DNA were detected (Figure 2G); nevertheless, as we have noted previously (46), the conformation of the homoduplex is likely not conducive to catalysis (see below).

FAN1 cleavage follows a similar pattern on CTG-harboring extrusions, with substrate preference in the following order: (CTG)_3_ (0.003 s^-1^) > (CTG)_2_ (0.0025 s^-1^) > (CTG)_4_ (0.0014 s^-1^) > (CTG)_5_ (0.001 s^-1^) > (CTG)_13_ (0.0007 s^-1^) > (CTG)_1_ (0.0006 s^-1^) (Figure 2C). It should be noted that a homoduplex DNA substrate is refractory to FAN1 cleavage (Supplementary Figure 2C) in line with our previously published work (39,46).

We then sought to determine the sites of FAN1 nuclease cleavage. In order to distinguish between the first endonuclease cut and subsequent incisions, we performed FAN1 nuclease assays using a (CAG)_2_ DNA substrate labeled on the 5’ end of the extrusion-containing strand. Denaturing PAGE sizing analysis of the products of FAN1 hydrolysis indicate that the primary endonuclease cleavage sites are located at positions 27 and 28 (1 and 2 nucleotides past the extrusion on the 3’ side, as measured from the 5’ end) (Figure 3A). We also observed a non-specific exonucleolytic cleavage from the 5’ DNA end on both DNA strands (Figure 3A, and Supplementary Figure 2D), an activity that was observed previously to yield a 2-4 nucleotide product (43). Time course experiments using a 3’ labeled DNA substrate revealed cleavage products at positions 18 and 19 as measured from the 3’ end (and in agreement with the 5’ labeled DNA measurements). We also observed subsequent nucleolytic cleavage sites at positions 15 and 13, followed by cuts after every nucleotide, with the cleavage mostly terminating upon removal of 10-12 nucleotides (Figure 2A). This suggests an exonucleolytic mechanism that ensues following early endonucleolytic incisions.

**Figure 3.**
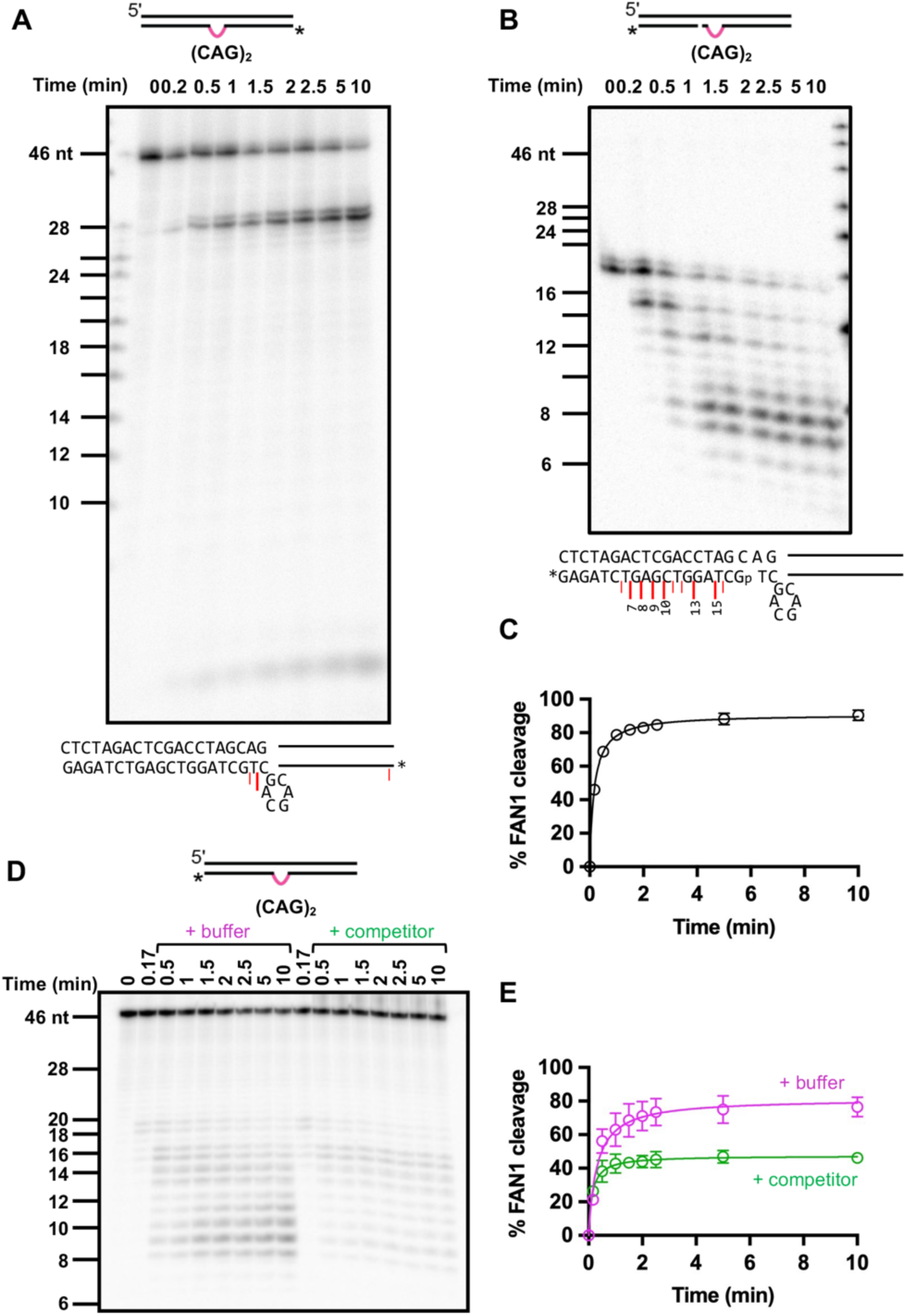
(**A**) Five nM of 5’-end radiolabeled DNA substrate harboring (CAG)_2_ extrusion was incubated with 11 nM FAN1 at 37 ⁰C in the presence of 5 mM MgCl_2_ and 70 mM KCl. Samples were collected at indicated time intervals and analyzed on 20% denaturing PAGE. The DNA marker is of the same sequence as the DNA substrate used in the experiment (see Supplementary Table 2). The image is a representative of n=2 independent experiments. The sequence of the DNA substrate with the major FAN1 endonucleolytic cleavage sites (red lines) is shown below the image. (**B**) A 3’-end radiolabeled DNA substrate harboring (CAG)_2_ extrusion and a single nick to mimic a DNA intermediate after the FAN1 endonuclease cleavage was used in this experiment; p indicates phosphorylation of the terminal nucleotide. Five nM of such DNA substrate was incubated with 11 nM FAN1, and samples were collected and analyzed as described in Figure 2A. The DNA marker is of the same sequence as the DNA substrate used in the experiment. (**C**) Quantification of gels as in (B). The graph represents mean values ± SD of n=3 independent experiments. (**D**) Ten nM of 3’-radiolabeled (CAG)_2_ DNA substrate was incubated with 2.5 nM FAN1 at 37 ⁰C in the presence of 5 mM MgCl_2_ and 70 mM KCl (see Materials and Methods). After 10 seconds of incubation at 37 ⁰C, either 250 nM Cy3-labelled (CAG)_2_ DNA substrate (competitor) or buffer was added to the reaction tubes (as indicated), and samples were collected at different time points as indicated. Cleavage products were resolved on 20% denaturing PAGE. (**E**) Quantification of experiments as in (D). The graph represents mean values ± SD of n=3 independent experiments.

In order to further understand the interdependence of the FAN1 endo– and exonuclease cleavage reactions, we prepared a DNA substrate that contains a (CAG)_2_ extrusion and a nick 2 nucleotides past the extrusion on the 3’ side to mimic the DNA intermediate post-endonucleolytic cleavage (Figure 3B, C). The observed initial rates of cleavage on this DNA substrate are 0.01 s-1 (∼2-fold faster than for the corresponding (CAG)_2_ substrate that does not harbor a nick, Figure 2B), suggesting that the initial incision by FAN1 might modestly limit the rate of the overall reaction.

Furthermore, as shown in Figure 3D, E, FAN1 nuclease cleavage of (CAG)_2_ extrusions occurs in a distributive manner, since challenge of the reaction with an excess of non-radiolabeled DNA substrate quenched the formation of additional cleavage products. This observed distributive behavior of the FAN1 nuclease on (CAG)_2_ extrusions has been reported previously for other DNA substrates (40,56).

### FAN1 cleaves (CAG)_2_ extrusions within CTG/CAG triplet repeat tracts

As mentioned above, extrahelical extrusions are formed due to strand slippage within repetitive sequences during cellular events that involve DNA helix opening and re-annealing. The experiments described above were focused on the effect of the DNA structure per se (size and sequence composition of the extrahelical extrusion) within a non-repetitive sequence context. To better mimic the natural context of the formation of CAG extrusions, we constructed a heteroduplex substrate in which the CAG extrusion is within a CTG/CAG repeat tract. Figure 4A, B, and Supplementary Figure 5 demonstrate that the heteroduplex (CTG)_11_/(CAG)_13_ (harboring two extra CAG repeats on one DNA strand) is an excellent substrate for the FAN1 nuclease. Unlike the substrates used in Figures 1 and 2, wherein the position of the extrusion is fixed within the overall sequence context, the extrusion within the (CTG)_11_/(CAG)_13_ sequence can form anywhere within the span of the repeat tract. While it is possible that the two extrahelical repeats exist as distinct and separate (CAG)_1_ extrusions, there is evidence suggesting that these extrahelical bases may be accommodated within a single (CAG)_2_ extrusion (57). Indeed, we observed the initial rates of FAN1 nuclease activity on this DNA substrate to be indistinguishable (∼0.005 s^-1^) from the DNA substrate harboring a (CAG)_2_ extrusion in the context of a non-repetitive DNA sequence (Figure 4A, B and Figure 2B). These data support the idea that the FAN1 nuclease targets the extrahelical extrusion itself rather than recognizing a specific sequence. However, unlike for the (CAG)_2_ substrate where we observed distinct cleavage sites (Figure 2A), the FAN1 cleavage on (CTG)_11_/(CAG)_13_ occurs throughout the DNA repeat tract (Figure 4A). This was also evident when we evaluated the FAN1 endonucleolytic cleavage using a 5’ labelled (CTG)_11_/(CAG)_13_ substrate: incisions occur throughout the CAG tract (Figure 4C, D) (unlike the (CAG)_2_ substrate where 2 distinct endonucleolytic cleavage sites were observed (Figure 2D)). The simplest explanation for this observation is that the (CTG)_11_/(CAG)_13_ substrate is composed of a population of DNA molecules wherein the CAG extrusion exists at different loci within the repetitive tract. It is also possible that the extrusion migrates within the repetitive tract (58). In either scenario, FAN1 activity on such DNA substrates would result in multiple cleavage products in this bulk measurement. FAN1 shows minimal cleavage on the fully paired homoduplex control containing (CTG)_13_/(CAG)_13_ repeat tract (Supplementary Figure 5B).

**Figure 4.**
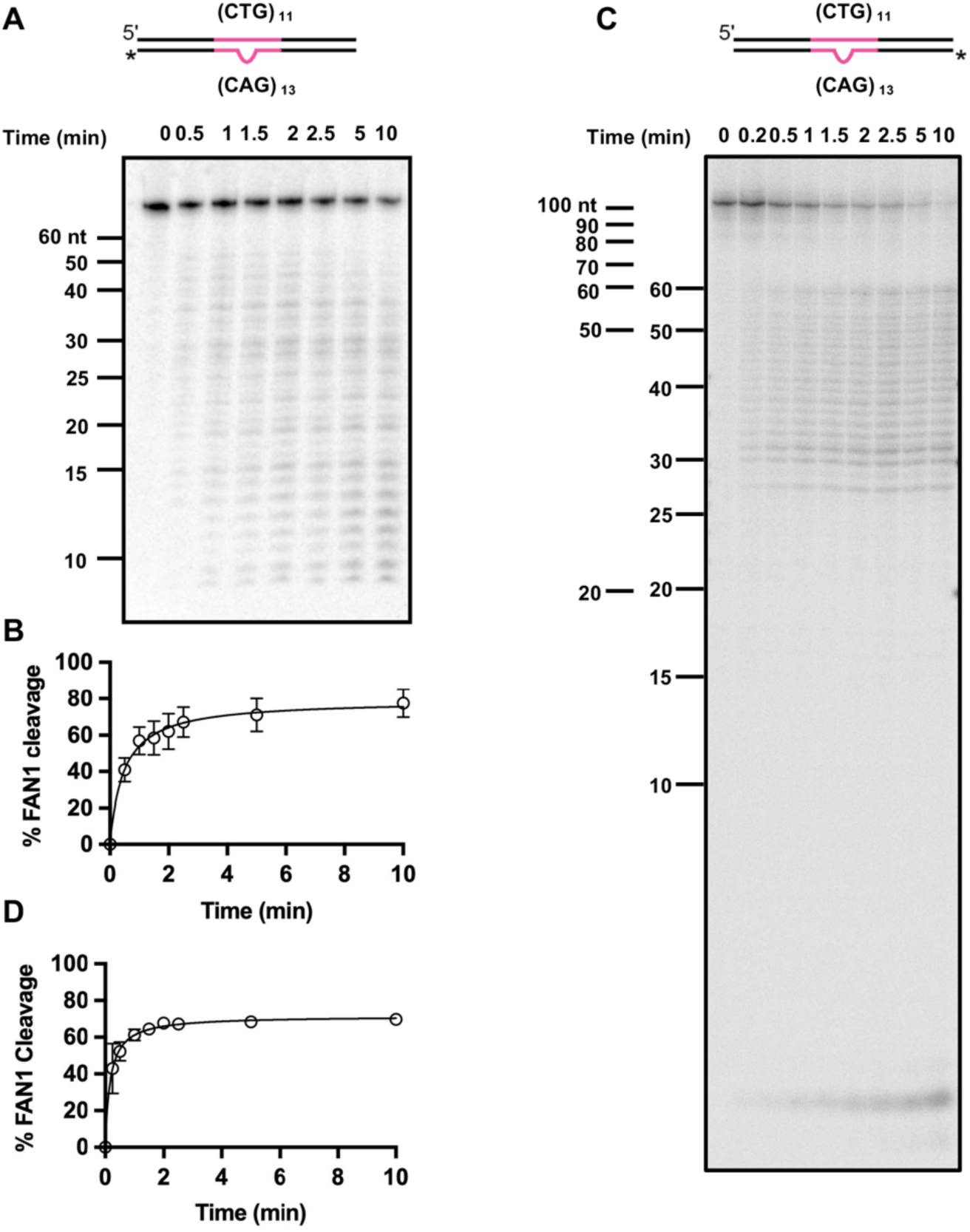
Flanking sequence of the extrusion does not affect FAN1 nuclease activity. (**A**) Five nM of 3’-radiolabeled (CTG)_11_/(CAG)_13_ DNA substrates were incubated with 11 nM FAN1 at 37 ⁰C in the presence of 5 mM MgCl_2_ and 70 mM KCl. Samples were collected at indicated time points and analyzed on 20% denaturing PAGE. The DNA size marker is shown. (**B**) Quantification of experiments as in (**A**). The graph represents mean values ± SD of n=3 independent experiments. (**C**) Experiment as in (**A**), except that a 5’-radiolabeled (CTG)_11_/(CAG)_13_ DNA substrate was used. The image is a representative of n=3 independent experiments with quantification in (**D**).

### DNA extrahelical extrusions induce DNA bending

Our FAN1 nuclease assays have revealed a strong preference of the FAN1 nuclease for (CAG)_2,3_ relative to (CAG)_1_ or homoduplex (Figures 1, 2, and 3). We hypothesized that these preferences may be governed by DNA conformational features unique to (CAG)_2,3_ versus (CAG)_1_. We therefore used AlphaFold (59) to model conformations adopted by these DNAs. As shown in Figure 5A, AlphaFold predicts that the extrahelical extrusion would induce a kink in the DNA with the degree of bend correlating with the size of the extrusion. Furthermore, two distinct classes of bent molecules are observed: extrusions containing C, CA, or (CAG)_1_ cause a modest bend of < 45°, whereas (CAG)_2_ and (CAG)_3_ result in a significant kink (> 90°) of the helical axis. Because bent DNAs display reduced electrophoretic mobility relative to corresponding linear DNAs (60–63), we experimentally tested the AlphaFold predictions by assessing whether extrahelical extrusions cause electrophoretic retardation. As shown in Figure 5B, our experiments reveal that first, electrophoretic retardation is a function of the location of the extrusion within the duplex such that extrusions located in the center of the molecule result in slower mobility relative to extrusions located closer to the DNA end. Since differential placement of the extrusion does not alter the molecular weight of these DNAs, effects on electrophoretic retardation may be attributed solely to the conformational features of these molecules. Second, we find that the size of the extrusion governs the extent of gel retardation, with the greatest degree of retardation observed for (CAG)_2_ and (CAG)_3_. Interestingly, extrusions containing C, CA, or (CAG)_1_ do not display appreciable gel retardation under these experimental conditions, consistent with the relative linearity of these molecules as per the AlphaFold predictions. Based on these findings, we propose that an extrusion-induced DNA bend of > 90° is a prerequisite for efficient recognition and cleavage by the FAN1 nuclease. This is in line with our recent cryo-EM structures of the PCNA-DNA-FAN1 complex wherein the FAN1 nuclease has an “L-shaped” conformation that can readily accommodate a (CAG)_2_ extrusion-harboring DNA that is bent by ∼100° (46).

**Figure 5.**
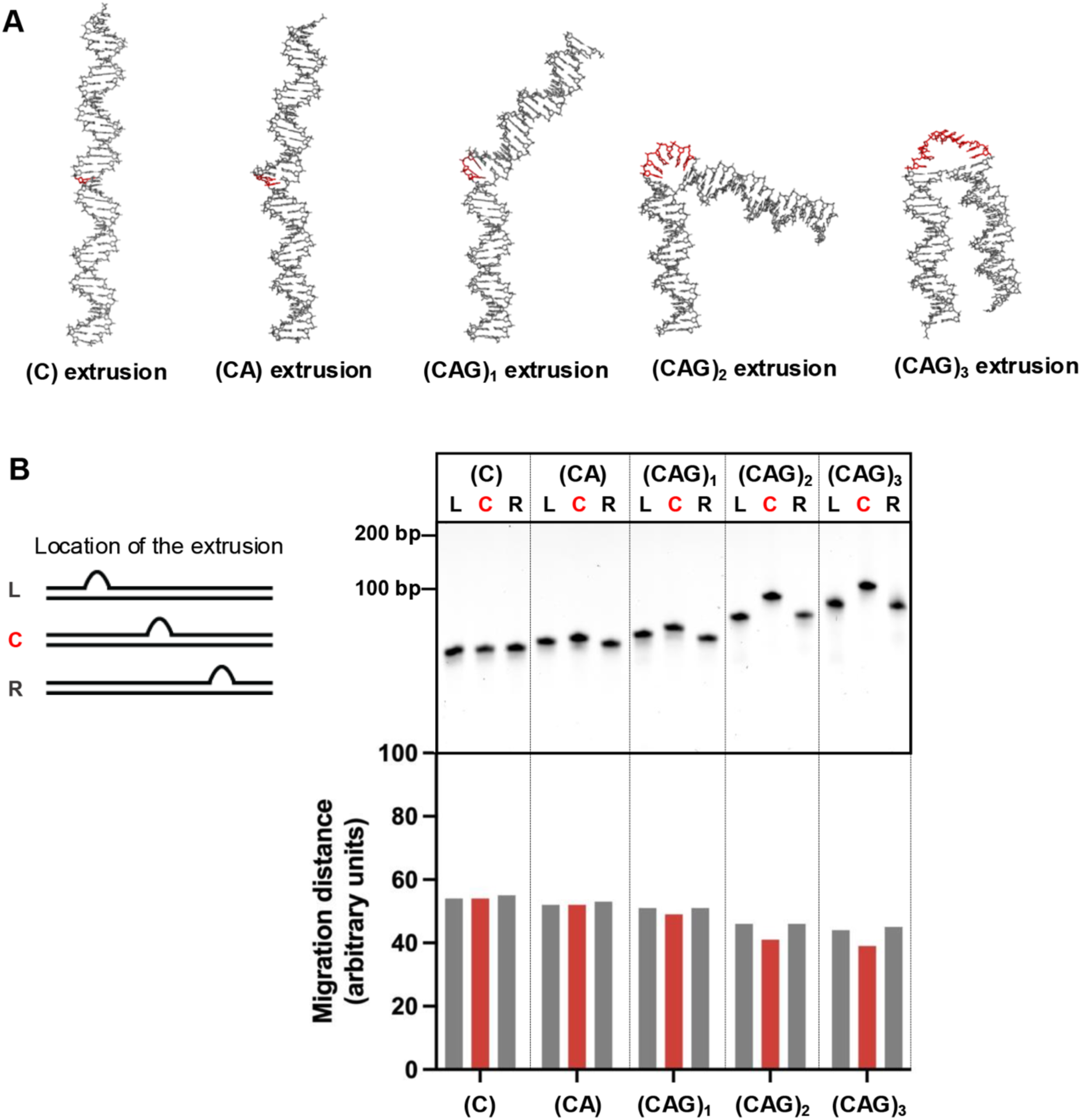
DNA extrahelical extrusions induce DNA bending. (**A**) AlphaFold structure prediction of DNA substrates harboring (C), (CA), (CAG)_1, 2, or 3_ extrahelical extrusions. (**B**) DNA substrates harboring indicated extrahelical extrusions either in the center (C), 10 base pairs away from a 5’ (L), or 3’ (R) DNA end were resolved on 20% PAGE. The migration distances from the top of the gel were measured, and quantifications are shown below. The experiment was repeated n=3 times with similar results. The oligonucleotide sequences used for constructing these DNA substrates are shown in Supplementary Table 3.

### Extrahelical extrusions are efficiently bound by MutSβ

A growing body of evidence suggests that triplet repeat expansions are controlled by two opposing pathways: whereas FAN1 prevents repeat expansions, MutSβ (a heterodimer of MSH2/MSH3 that recognizes insertion/deletion loops (18)) drives this process (21,32,33). Our recent studies have shown that FAN1 can compete with MutSβ for binding to (CAG)_2_ extrahelical extrusions, and in the absence of MutSβ can efficiently mediate their removal (39). We have therefore proposed that this competition for CAG extrusion binding may underly pathway choice between MutSβ-driven CAG expansion, and FAN1-dependent CAG stabilization. To further dissect the molecular basis of this pathway choice, we asked whether MutSβ (like FAN1) has a binding preference for (CAG)_2_ over (CAG)_1_. SPRS sensorgrams for time-dependent association and dissociation of MutSβ with (CAG)_1_, (CAG)_2,_ (CAG)_3_, (CAG)_13,_ and a homoduplex control (Figure 6) were analyzed by global fit to a 1:1 binding model (according to Eq (3) in Materials and Methods). The on rate/off rate correlation plot (Figure 6C), and steady-state isotherm (Figure 6D) established that (i) binding of MutSβ to (CAG)_1_, (CAG)_2,_ and (CAG)_3_ is significantly tighter than to the homoduplex control (in agreement with our previous findings (35)), and (ii) MutSβ shows no significant preference for (CAG)_2_ over (CAG)_1_ or (CAG)_3_. Further, MutSβ displays a ∼10-fold higher affinity for (CAG)_1_, (CAG)_2_, and ∼ 5-fold higher for (CAG)_3_ over (CAG)_13_ with a K_D_ of 0.54 ± 0.1 nM, 0.5 ± 0.16 nM, 0.97 ± 0.2 nM, and 5.09 ± 0.05 nM, respectively.

**Figure 6.**
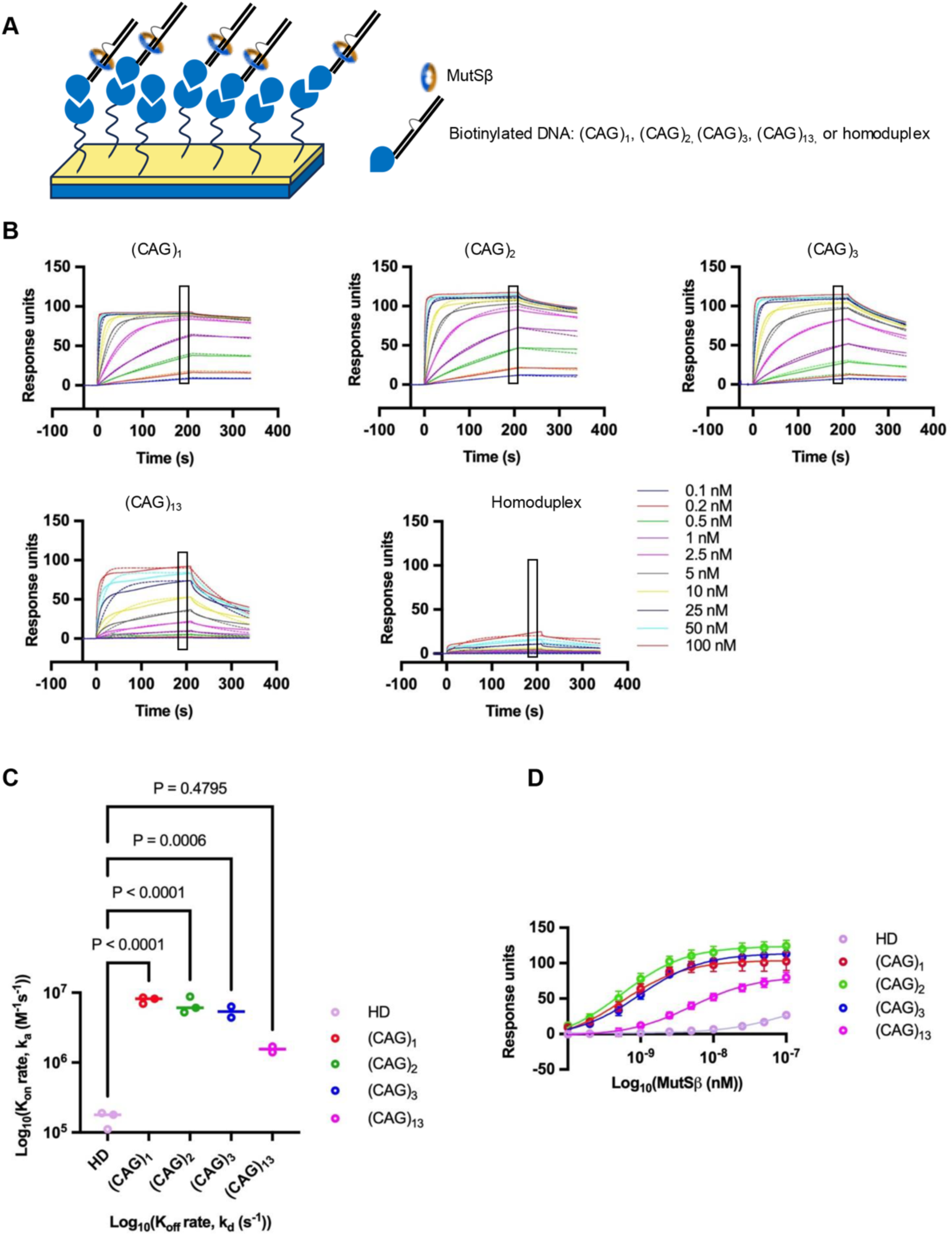
Extrahelical extrusions are efficiently bound by MutSβ. (**A**) Schematics of MutSβ-DNA interaction on the streptavidin sensor chip. (**B**) Binding sensograms for homoduplex, (CAG)_1_, (CAG)_2_, (CAG)_3_, and (CAG)_13_ with MutSβ concentration from 0.1 nM to 100 nM is shown by solid lines and the dashed lines corresponds to the kinetic fitting for 1:1 binding model, using Eq (3) (Materials and Methods). (**C**) Plot for on-off chart displays association (k_on_) and dissociation (k_off_) rates calculated from kinetic fitting data for (CAG)_1_, (CAG)_2_ (CAG)_3_, (CAG)_13_ and homoduplex substrates, obtained from 3 independent experiments ((CAG)_3_ and (CAG)_13_ data are based on n=2 independent experiments) with error bars representing SD. (**D**) Steady-state affinity plots for homoduplex, (CAG)_1_, (CAG)_2_, (CAG)_3_, and (CAG)_13_ from affinity fitting using Eq. (4) (Materials and Methods).

### Substrate preferences for PCNA-RFC-dependent FAN1 and MutSβ-dependent MutLα nuclease reactions

The physical and functional interactions between DNA and MutSβ or FAN1 described above were evaluated in simple binary reactions. However, we have previously shown that at physiological ionic strength, activation of the FAN1 nuclease on circular heteroduplexes harboring a (CAG)_2_ extrusion requires interaction between FAN1 and DNA-loaded PCNA (a process catalyzed by the RFC clamp loader) (39,46). For these experiments we used a circular DNA harboring a nick to enable efficient PCNA loading by RFC. As shown in Figure 7A, PCNA-activated FAN1 efficiently cleaves DNAs harboring (CAG)_2_ or (CAG)_3_ (appearance of the faster migrating band), but not a (CAG)_1_ extrusion (no detectable cleavage on the complementary DNA strand was observed (Figure 7C)). Thus, the preference for (CAG)_2_ or (CAG)_3_ over (CAG)_1_ is an intrinsic feature of the FAN1 nuclease.

**Figure 7.**
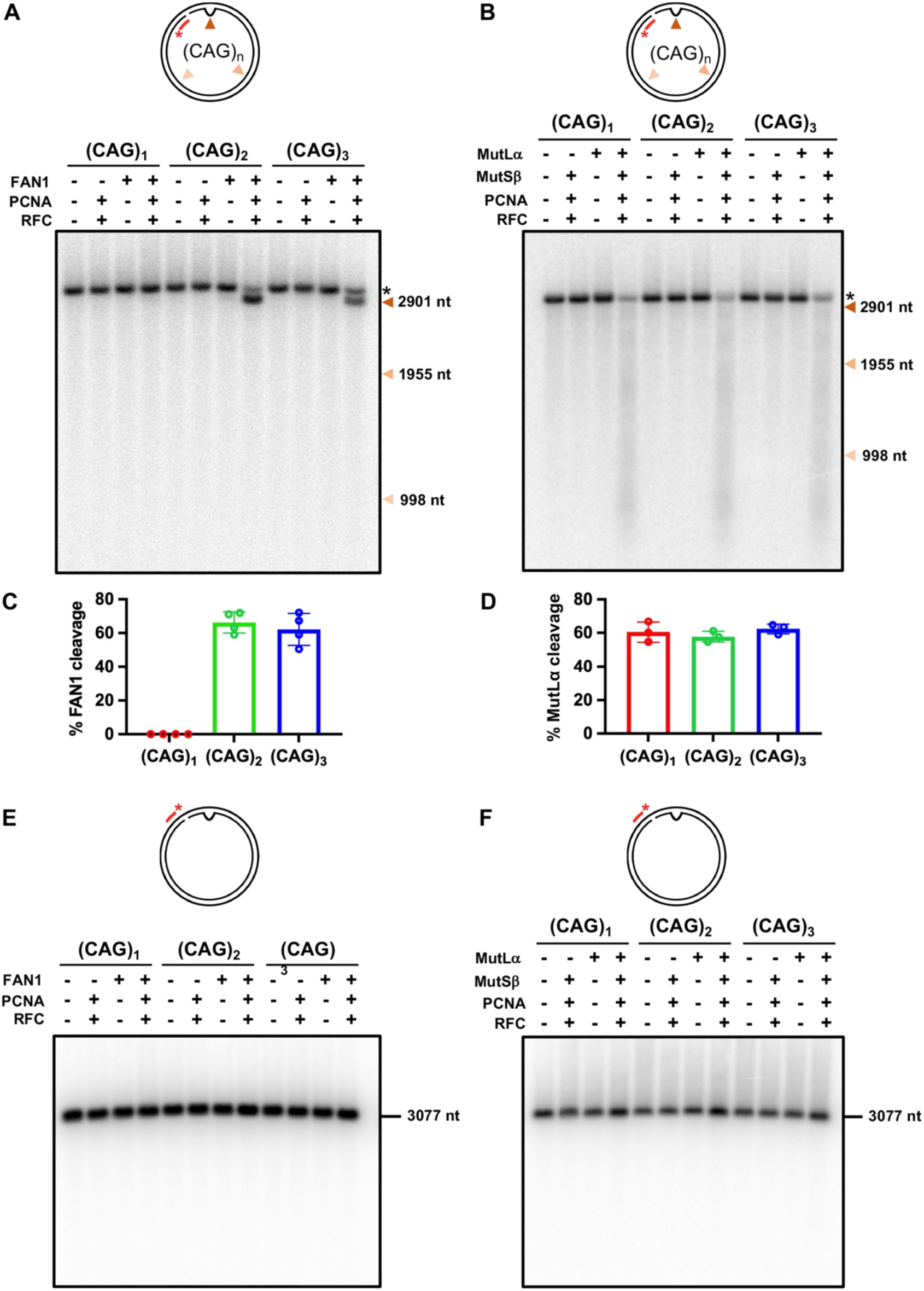
Substrate preferences for PCNA-RFC-dependent FAN1 and MutSβ-dependent MutLα nuclease reactions. (**A**) Circular DNA substrates harboring (CAG)_1_, (CAG)_2_, or (CAG)_3_ extrusion and a single nick located 3’ to the extrusion were incubated with FAN1, PCNA and RFC (as indicated) in a buffer containing 5 mM MgCl_2_ and 125 mM KCl (Materials and Methods) for 10 min at 37 °C. Products of the reaction were resolved on 1% denaturing agarose gels, followed by hybridization with ^32^P-labeled oligonucleotide probe (Fwd 2943). The red arrow indicates the location of the extrusion; the black asterisk indicates a full-length substrate. Size markers are shown. The graph below the image represents percent of FAN1 cleavage in the complete reaction (in the presence of PCNA and RFC). Graph represents mean values ± SD of n=4 independent experiments. (**B**) Substrates as in (A) were incubated with MutSβ, MutLα, PCNA, and RFC (as indicated) in a buffer containing 5 mM MgCl_2_ and 125 mM KCl (Materials and Methods) for 15 min at 37 °C. Products of the reactions were analyzed as in (A). Black asterisk indicates a full-length substrate. The graph below the image represents the percent of MutLα cleavage in the complete reaction (in the presence of MutSβ, PCNA and RFC). The graph represents mean values ± SD of n=3 independent experiments. (**C, D**) Products of the reaction from A and B, respectively, were resolved on 1% denaturing agarose gels, followed by hybridization with ^32^P-labeled oligonucleotide probe (Rev 2970) to visualize the complementary DNA strands. The images are representative of n=2 independent experiments.

By contrast, MutSβ did not display binding preference between (CAG)_1_, (CAG)_2_, and (CAG)_3_ (Figure 6). Because extrusion-bound MutSβ initiates DNA mismatch repair by activating the MutLα endonuclease on nicked circular heteroduplexes in a RFC and PCNA-dependent manner (38,39), we used this reaction as an indicator of initiation of MutSβ-dependent repair of (CAG)_1_, (CAG)_2_, and (CAG)_3_ extrusions. Unlike FAN1, MutSβ-dependent MutLα endonuclease activity occurred with comparable facility on all three extrusions (appearance of a smear) (Figure 7B). No cleavage was detected on the complementary DNA strand (Figure 7D). Thus, taken together with the FAN1 nuclease and DNA binding assays, these results suggest that DNA substrates harboring a (CAG)_1_ extrusion may be exclusively processed by a MutSβ-dependent pathway, whereas (CAG)_2, 3_ and (CAG)_13_ are likely subject to competition between FAN1 and MutSβ.

## DISCUSSION

Triplet repeat expansion is a net consequence of the interplay between opposing pathways: whereas MutSβ initiated mismatch repair drives repeat expansion, a FAN1-dependent mechanism prevents this process (21,32,33). We have demonstrated that a plausible molecular explanation for the role of FAN1 in attenuating repeat expansion is the ability of the FAN1 nuclease to remove extrahelical extrusions formed within triplet repeat tracts via a pathway that relies on short-patch DNA excision/resynthesis (39). Because extrahelical extrusions can also be processed by a MutSβ-dependent pathway (35,39), we have proposed that the competition between FAN1 and MutSβ for such extrusions could determine the choice between repeat stability versus instability. Indeed, relative molar stoichiometries of FAN1 and MutSβ are key determinants that drive extrusion processing in vitro (39). However, DNA strand mishybridization within repetitive sequences can lead to the formation of a variety of extrahelical extrusions and hairpin-loop structures (2–4,64–66). Could it be that the size of the extrusion plays a role in dictating pathway choice?

Therefore, we sought to determine the preferred DNA extrahelical extrusion sizes for processing by FAN1 versus MutSβ. Interestingly, we observed that whereas extrusions of 2-3 repeats efficiently provoke FAN1 nuclease activity, extrusions of one triplet repeat are refractory to cleavage by FAN1. This nucleolytic cleavage preference can be explained (at least in part) by the preferential binding of FAN1 to these structures: FAN1 binds to (CAG)_2, 3_ extrusions efficiently, but very poorly to (CAG)_1_ extrusions and does not reach saturation (Figure 2G, H). By contrast, MutSβ binding is largely agnostic to the size of the extrahelical extrusion; the protein binds with a similar high affinity to extrusions composed of one, two, or three repeats (Figure 6). We have previously shown that at physiological ionic strength, FAN1 is activated by the presence of DNA-loaded PCNA on a DNA substrate harboring a (CAG)_2_ or (CTG)_2_ extrusion (Figure 7A and (39)). Under these conditions, (CAG)_1_ is refractory to PCNA-activated FAN1 cleavage (Figure 7A). By contrast, MutSβ-dependent MutLα endonuclease in the presence of DNA-loaded PCNA processes (CAG)_1_, (CAG)_2_, and (CAG)_3_ with comparable facility. Thus, the overwhelming preference of FAN1 for extrusions containing two or three repeats (over a one-repeat extrusion) is consistent across nuclease assay conditions and binding studies. Likewise, the lack of preference of MutSβ for (CAG)_1, 2, or 3_ is evident in binding studies as well as MutSβ-dependent MutLα nuclease assays. These observations highlight the fact that extrusion size could be a key driver of pathway choice. Interestingly, (CAG)_13_ extrusions that have been proposed as hairpin intermediates in the expansion process are not only poor substrates for FAN1 binding and nucleolytic cleavage in our in vitro studies, but also are bound by MutSβ with a ∼5 to 10-fold lower affinity than (CAG)_1_, (CAG)_2,_ and (CAG)_3_. It is noteworthy in this regard that triplet repeat expansions in iPSC-derived striatal neurons and in animal models occur in 1 repeat increments (12,21,67). Therefore, the role of such hairpin structures in driving pathway choice is unclear.

What might be the molecular basis of the FAN1 binding specificities reported here? A clue comes from our recent cryo-EM studies of the FAN1-PCNA-DNA ternary complex wherein the DNA is bent at an outer angle of ∼102° at the (CAG)_2_ extrusion site, thus positioning the extrusion appropriately for catalysis (46). Given that the overall architecture of the FAN1 protein tracks with the DNA bend (46,56), we hypothesize that the bend angle of the DNA is a key determinant that facilitates efficient FAN1 catalysis. AlphaFold modelling and gel retardation studies indicate that (CAG)_1_ extrusions induce a very modest kink (∼40°) in the DNA helix, as compared with a significant (>90°) kink for (CAG)_2_ or (CAG)_3_. Thus, the intrinsic bendability of the DNA at the extrusion site likely facilitates cleavage by FAN1. Indeed, the crystal structure of FAN1 bound to its *bona fide* double fork substrate is characterized by a DNA outer angle bend of ∼104° at the strand break (56). Since FAN1 can bind to homoduplex DNA to an appreciable degree at sufficiently high concentrations (46), why such molecules are nevertheless refractory to FAN1 cleavage has remained a question. This resistance to cleavage by FAN1 could be explained by the lack of an intrinsic bend in homoduplex DNAs. It is therefore tempting to speculate that FAN1 may be a general sensor of DNA lesions as well as unusual DNA conformations that produce a strong kink in the DNA duplex. We are currently investigating these questions.

The overarching goal of our investigations is to enhance our understanding of the protein and nucleic acid factors that drive DNA repeat expansions in human disease. Our observations lend themselves to a model wherein (CAG)_1_ extrusions formed by strand slippage within long CAG/CTG repeats are preferentially processed by MutSβ-dependent pathway, thereby leading to expansion. On the other hand, extrusions containing two or three repeats can be processed by either pathway depending on relative protein stoichiometry. Thus, under cellular conditions of spatio-temporal abundance of MutSβ-associated DNA mismatch repair proteins, (CAG)_2_ and (CAG)_3_ extrusions would provoke an expansion-prone repair process. However, processing of such extrusions when FAN1 is predominant would result in stabilization of the repeated tract. This complex interplay between MMR and FAN1 could also be modulated by a physical interaction between FAN1 and MLH1, a topic that we are actively pursuing.

## DATA AVAILABILITY

All data pertaining to this article are available in the article and in its supplementary materials.

## SUPPLEMENTARY DATA

Supplementary Data are available as a separate PDF file.

## AUTHOR CONTRIBUTIONS

Mayuri Bhatia: Conceptualization, Formal analysis, Methodology, Validation, Writing—original draft & editing. Ashutosh Phadte: Methodology. Anna Lakhina: Methodology. Anthony Monte Carlo III: Methodology. Sarah Barndt: Methodology. Anna Pluciennik: Conceptualization, Formal analysis, Writing—review & editing.

## ACKNOWLEDGEMENTS

We thank Dr. Paul Modrich (Duke University Medical Center) for sharing the DNA constructs to produce circular DNA substrates, and Ravi R. Iyer for helpful discussions.

## SUPLEMENTARY MATERIALS

### Supplementary Figure 1-5

**Supplementary Figure 1.**
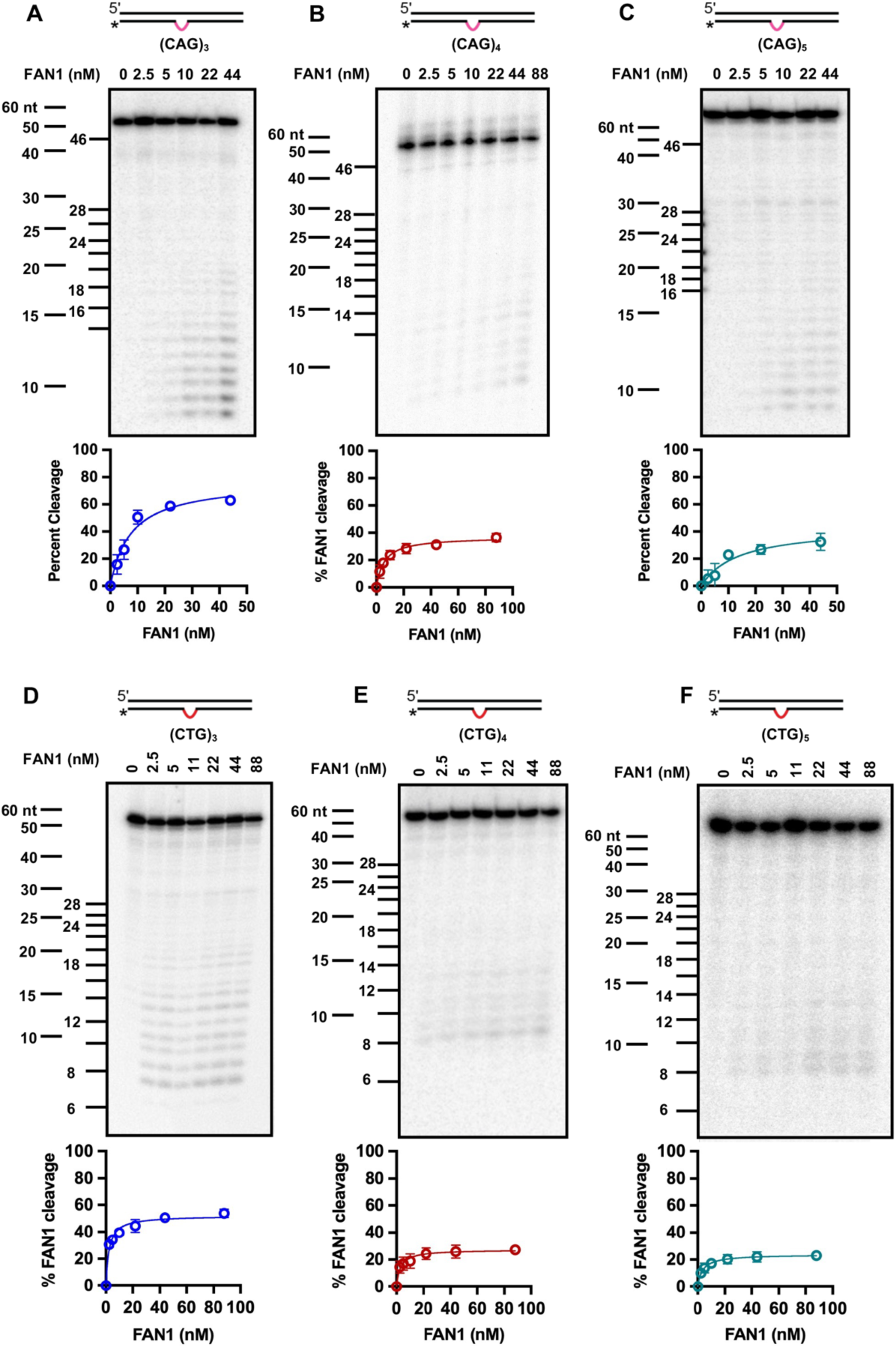
FAN1 preferentially cleaves extrahelical extrusions containing 6-9 nucleotides. 3’-end radiolabeled DNA substrates harboring (**A**) (CAG)_3_, (**B**) (CAG)_4_, (**C**) (CAG)_5_, (**D**) (CTG)_3_, (**E**) (CTG)_4_, (**F**) (CTG)_5_ were incubated with increasing concentrations of FAN1 for 10 min (see Materials and Methods). Samples were collected at indicated time points and resolved on 20% denaturing PAGE. Representative images are shown. Quantification of the percentage of FAN1 cleavage is shown below each respective image. Graphs are presented as mean values ± SD of at least 3 independent experiments.

**Supplementary Figure 2.**
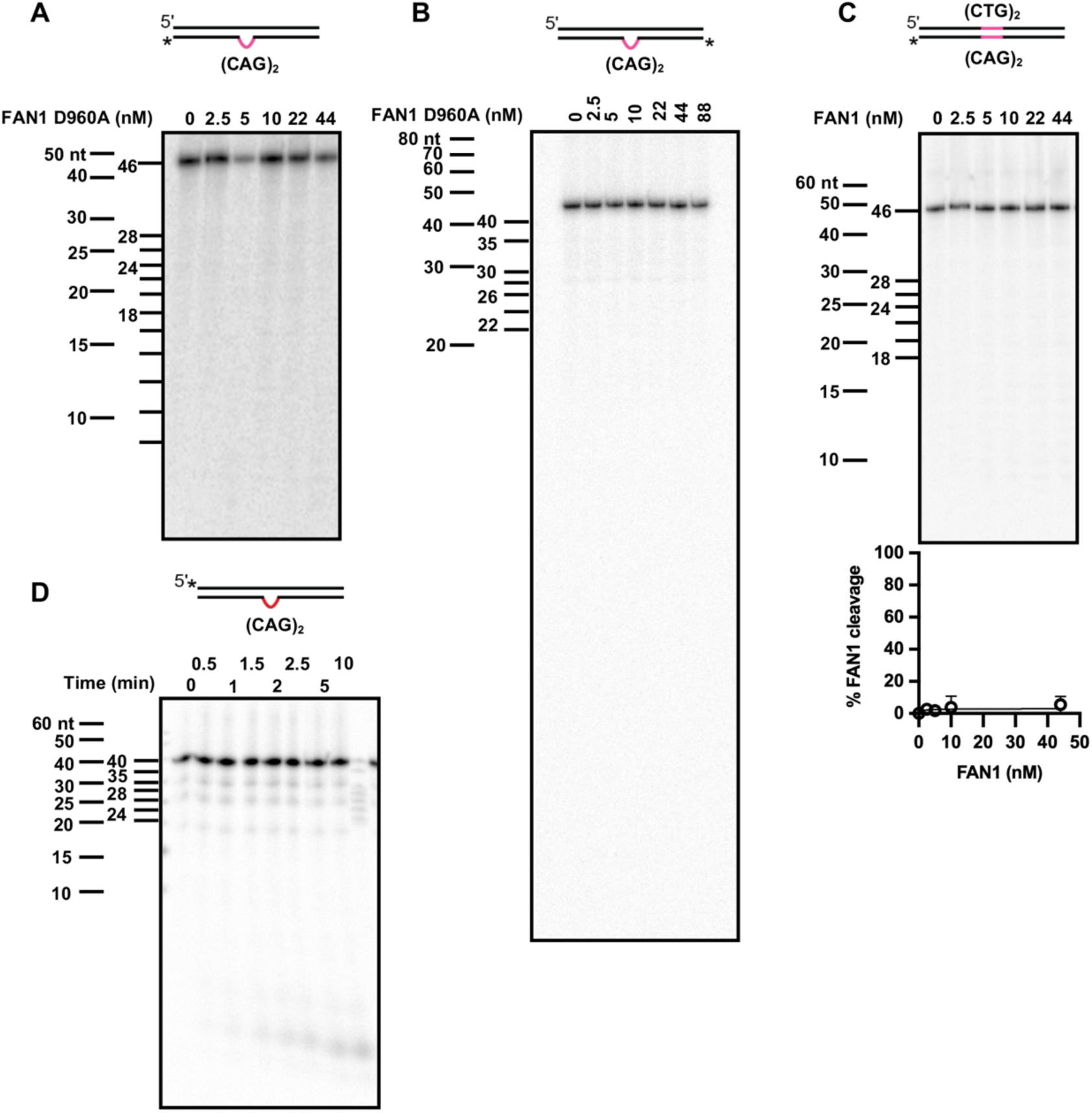
(**A, B**) Five nM 3’-end radiolabeled or 5’-end radiolabeled (CAG)_2_ DNA substrate, respectively, was incubated with increasing concentrations of a catalytically dead mutant of FAN1 (D960A) at 37 ⁰C in the presence of 70 mM KCl and 5 mM MgCl_2_ for 10 min. Samples were collected and resolved on 20% denaturing PAGE. The images are representative of n=2 independent experiments. (**C**) Five nM of 3’-radiolabeled homoduplex DNA was incubated with 11 nM FAN1 at 37⁰C in the presence of 70 mM KCl and 5 mM MgCl_2_. Samples were collected and analyzed as in (**A**). The image is a representative of n=3 independent experiments. (**D**) Five nM DNA substrate harboring (CAG)_2_ extrusion (that was 5’-radiolabeled on the complementary DNA strand) was incubated with 11 nM of FAN1 at 37 ⁰C for for indicated time in the presence of 70 mM KCl and 5 mM MgCl_2_. Samples were collected and analyzed as in (**A**). The image is a representative of n=2 independent experiments.

**Supplementary Figure 3.**
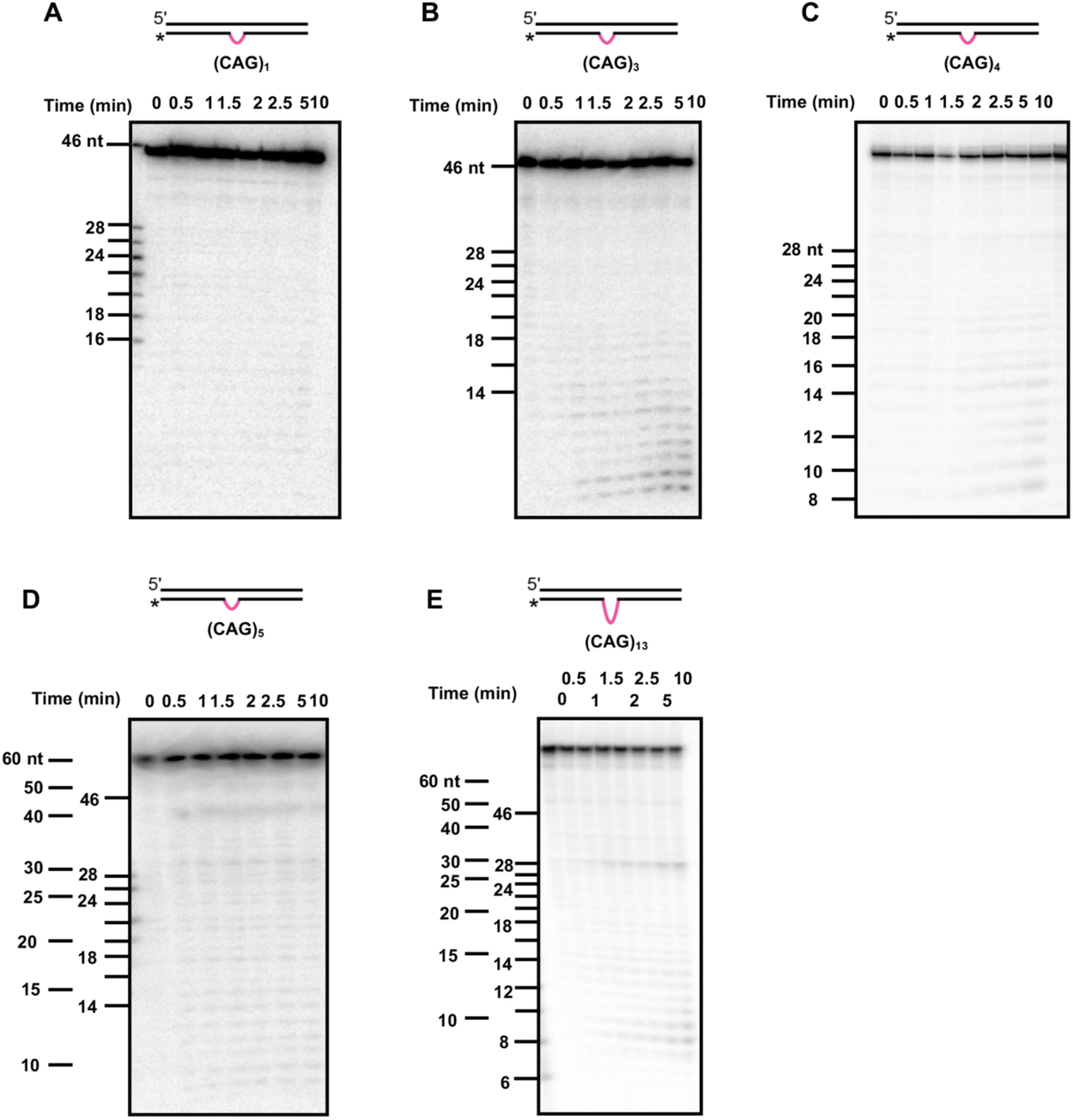
FAN1 cleavage rates depend on the size of the CAG extrusion. Five nM of 3’-radiolabeled DNA substrates harboring (**A**) (CAG)_1_, (**B**) (CAG)_3_, (**C**) (CAG)_4_, (**D**) (CAG)_5_, (**E**) (CAG)_13_ were incubated with 11 nM FAN1 at 37 ⁰C in presence of 70 mM KCl and 5 mM MgCl_2_. Samples were collected at indicated time points and resolved on 20% denaturing PAGE. The images are representatives of n=3 independent experiments (except for (CAG)_4_ where the experiment was repeated two times).

**Supplementary Figure 4.**
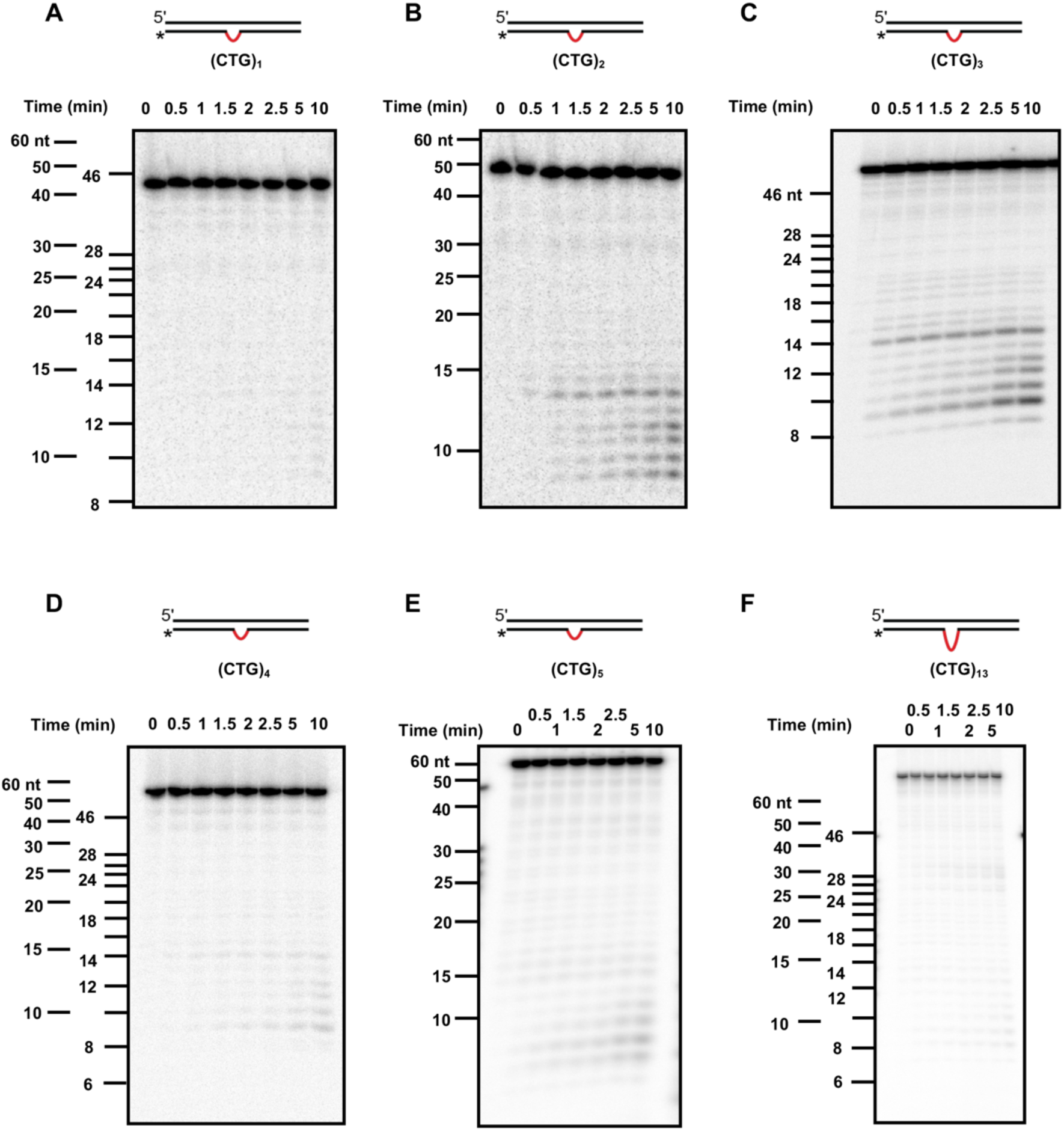
FAN1 cleavage rates depend on the size of the CTG extrusion. Five nM of 3’-radiolabeled DNA substrates harboring (**A**) (CTG)_1_, (**B**) (CTG)_2_, (**C**) (CTG)_3_, (**D**) (CTG)_4_, (**E**) (CTG)_5_, (**F**) (CTG)_13_ were incubated with 11 nM FAN1 at 37 ⁰C in the presence of 70 mM KCl and 5 mM MgCl_2_. Samples were collected at indicated time points and resolved on 20% denaturing PAGE. The image is a representative of n=3 independent experiments (except for (CTG)_3_, the experiments were repeated two times).

**Supplementary Figure 5.**
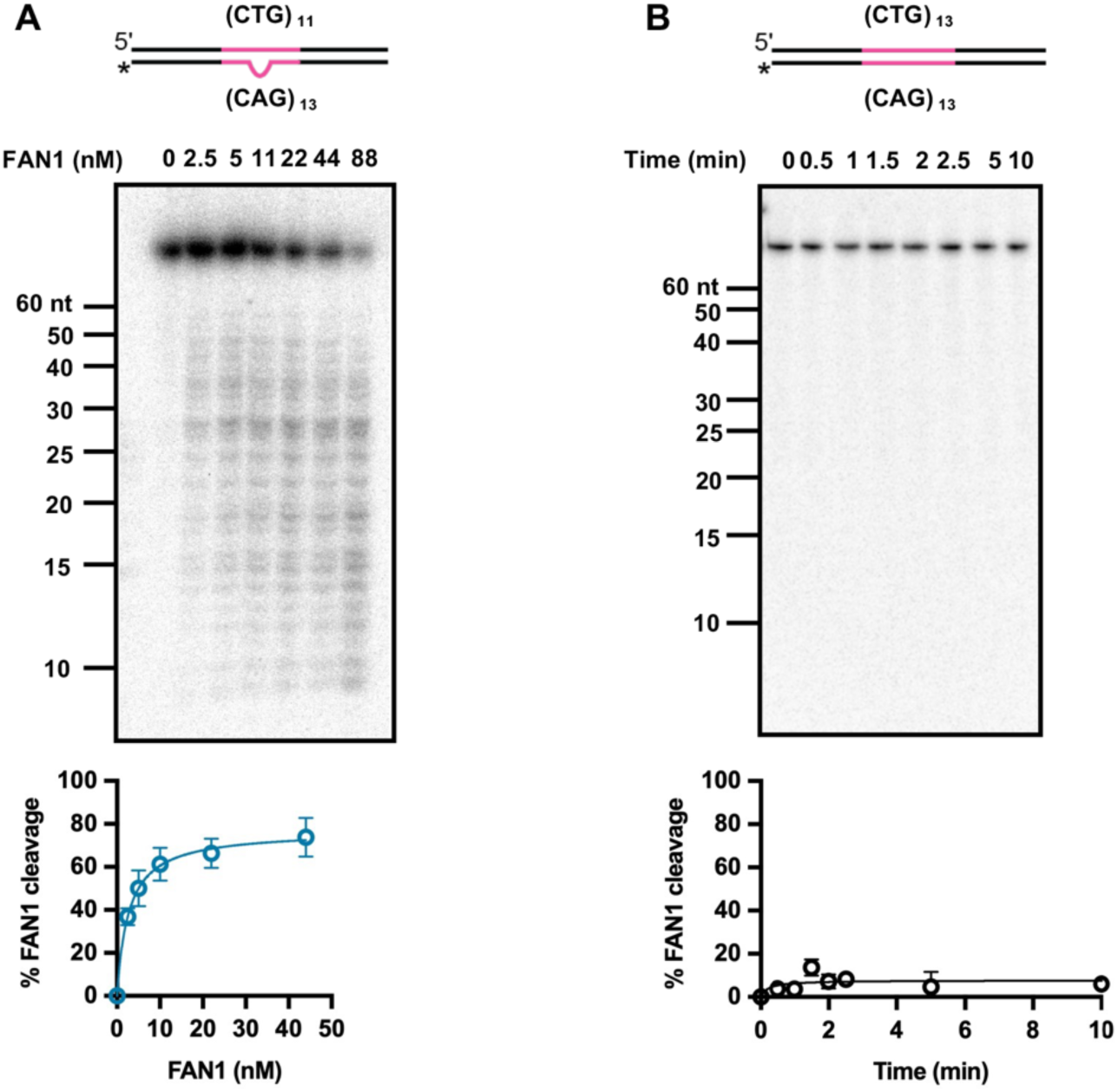
(**A**) Five nM 3’-end radiolabeled (CTG)_11_/(CAG)_13_ DNA substrate was incubated with increasing concentrations of FAN1 at 37 ⁰C in a buffer containing 70 mM KCl and 5 mM MgCl_2_ for 10 min. Samples were collected and resolved on 20% denaturing PAGE. Quantification of percent of FAN1 cleavage is shown below. Graph represents mean values ± SD of n=5 independent experiments. (**B**) Five nM 3’-end radiolabeled (CTG)_13_/(CAG)_13_ homoduplex DNA was incubated with 11 nM FAN1 at 37 ⁰C in presence of 70 mM KCl and 5 mM MgCl_2_. Samples were collected at indicated time points and analyzed as in (**A**). Quantification of percent of FAN1 cleavage is shown below. Graph represents mean values ± SD of n=3 independent experiments.

